# Low doses of radiation increase the immunosuppressive profile of lung macrophages during viral infection and pneumonia

**DOI:** 10.1101/2020.05.11.077651

**Authors:** Lydia Meziani, Charlotte Robert, Marion Classe, Bruno Da Costa, Michele Mondini, Céline Clemenson, Pierre Mordant, Samy Ammari, Ronan Le Goffic, Eric Deutsch

## Abstract

Severe pneumonia and acute respiratory distress syndrome (ARDS) have been described in patients with severe COVID-19. Recently, early clinical data reported the efficacy of low doses of radiation therapy (RT) in the treatment of ARDS in patients with severe COVID-19. However, the involved mechanisms remained unknown. Here, we used airways-instilled lipopolysaccharide (LPS) and influenza virus (H1N1) as murine models of pneumonia, and Tolllike receptor (TLR)-3 stimulation in human lung macrophages. Low doses RT (0.5-1 Gy) decreased LPS induced pneumonia, and increased the percentage of Nerve- and Airway-associated Macrophages (NAMs) producing IL-10. During H1N1 viral infection, we observed decreased lung tissue damage and immune cell infiltration in irradiated animals. Low doses RT increased IL-10 production by infiltrating immune cells into the lung. Irradiation of TLR-3 ligand-stimulated human lung macrophages *ex vivo* increased IL-10 secretion and decreased IFNγ production in the culture supernatant. The percentage of human lung macrophages producing IL-6 was also decreased. Our data highlight one of the mechanisms by which low doses RT regulate lung inflammation and skew lung macrophages towards an anti-inflammatory profile. These data provide the preclinical rationale for the use and for the optimization of low doses RT in situations such as COVID-19-induced ARDS.

## Introduction

The COVID-19 pandemic is responsible for more than one million deaths and 37.39 million cases worldwide as of October 12, 2020. The responsible agent, SARS-CoV-2, is an enveloped RNA virus of the Coronaviridae virus family. Human-to-human transmission occurs through respiratory droplets or contaminated surfaces (1). The average incubation period is 5 days, with a range of 1 to 14 days. Most patients present mild respiratory tract infections, most commonly characterized by fever (82%) and cough (81%). Severe pneumonia and acute respiratory distress syndrome (ARDS) have been described in 14% of the reported cases, and the overall mortality is around 1-2% (2). Current therapeutic approaches involve mechanical ventilation, acute supportive care, management of organ failure, steroids and antiviral therapies such as remdesivir, lopinavir–ritonavir and interferon beta-1, dexamethasone, which are currently under investigation. Growing information suggests that patients with severe COVID-19 have a marked inflammatory state characterized by a cytokine storm syndrome similar to that seen in severe influenza cases (3), and high levels of both calprotectin and myelopoiesis (4).

Chest irradiation at low dose has been used successfully in the past to treat pneumonia, especially before the onset of effective antimicrobial agents. The estimation of the radiation doses absorbed by the lungs during orthovoltage irradiation techniques rendered complex at that time. Clinical reports suggest an early improvement of breathing difficulties within hours and a reduction of mortality (5–7). Currently, several clinical trials are underway (referenced on clinicaltrial.gov) and some studies, even if in small cohorts, confirmed the efficacy of low doses of RT in the treatment of ARDS in patients with severe COVID-19 (8–10). However, the involved mechanisms remain unknown and the use of chest radiation therapy (RT) for COVID patients has been the subject of a vivid scientific controversy. Amongst the authors, for and against arguments, is the absence of proof of concept as well as the intrinsic risk that radiation induces lung damage and boosts viral expansion after RT.

RT exerts well-known anti-inflammatory properties when used at doses up to 1Gy while producing pro-inflammatory effects at higher doses (11), highlighting the complexity of the immunological mechanisms and the interrelationship between ionizing radiation (IR) and inflammation.

Pulmonary macrophages have been implicated in maintaining lung homeostasis by immune surveillance and clearance of dead cells, debris, and invading pathogens. The lung harbors two distinct populations of macrophages, alveolar macrophages (AMs) and interstitial macrophages (IMs) (12, 13). AMs are located in alveolar space and seem to play a direct antiviral role since AM depletion yields higher viral loads. IMs are located in the interstitium, along with dendritic cells and lymphocytes. Recently, a new IM subpopulation of Nerve- and Airway-associated Macrophages (NAMs), have been characterized in mice (14). NAMs are distinct from other lung-resident macrophage subsets and highly express immunoregulatory genes. NAMs proliferate robustly after influenza infection and activation with the polyinosinic:polycytidylic acid Poly(I:C), and in their absence the inflammatory response is augmented, resulting in excessive production of inflammatory cytokines and innate immune cell infiltration. NAMs function is to maintain immune and tissue homeostasis, and regulate infection-induced inflammation through the secretion of immunosuppressive factors such as IL-10. Viral infection in the absence of NAMs is associated with an excess of proinflammatory cytokines and chemokines such as IL-6, CCL2, CCL3, and CCL5 eventually leading to massive lung damage and death (14). Interestingly, potential pathological roles of macrophages during SARS-CoV-2 infection have been described (15).

Irradiation has a direct effect on macrophage activation, depending on time, dose, and subpopulations. In the tumor stroma, high doses of IR (> 8Gy) promote anti-inflammatory activity of macrophages (16) and low doses (< 2Gy) alone or combined with immunotherapy induce proinflammatory activity of macrophages (17, 18). Within the lung, irradiation at high dose (16Gy) affects the phenotype of alveolar and interstitial macrophages differently, resulting in distinct local cytokine and chemokine microenvironments in the tissue and alveoli. Interestingly, we have observed a difference between lung sub compartments in response to fibrogenic irradiation dose (16Gy) with an immune response first in the parenchyma, then in the alveolar compartment (19).

In the present study, using Lipopolysaccharide (LPS), influenza A PR8 virus (H1N1) or Toll-like receptor 3 (TLR3) ligand Poly(I:C) as inductors of inflammation (20), and low doses of RT as a potential treatment, we investigated whether RT could be involved in counteracting lung inflammation through IL-10 production.

## Results

### Low doses of RT protect mice from LPS-induced pneumonia and increase IL-10 production by NAMs

To investigate whether low doses of RT protect mice from lung inflammation, we treated mice with sublethal doses of LPS by intratracheal administration over two consecutive days. Six hours after the second intratracheal administration, mice were irradiated at the whole thorax at 0.5 or 1 Gy and underwent Computed tomography (CT) imaging at lung level at several time points after whole thorax irradiation. LPS administration significantly increased lung density at 96 hours (post administration of the first dose of LPS) compared to PBS control, suggesting the development of an inflammatory process in the lungs (Fig. 1A). Interestingly, irradiated lungs at 1Gy had a reduced tissue density compared to the LPS group, suggesting that a low dose of 1 Gy protects mice from LPS-induced lung inflammation. Furthermore, histopathological analysis showed a decrease in parenchyma destruction (Fig. 1B) and a trend towards a reduced peribronchial infiltration in irradiated lungs at 1Gy (Fig. 1C).

**Fig. 1.**
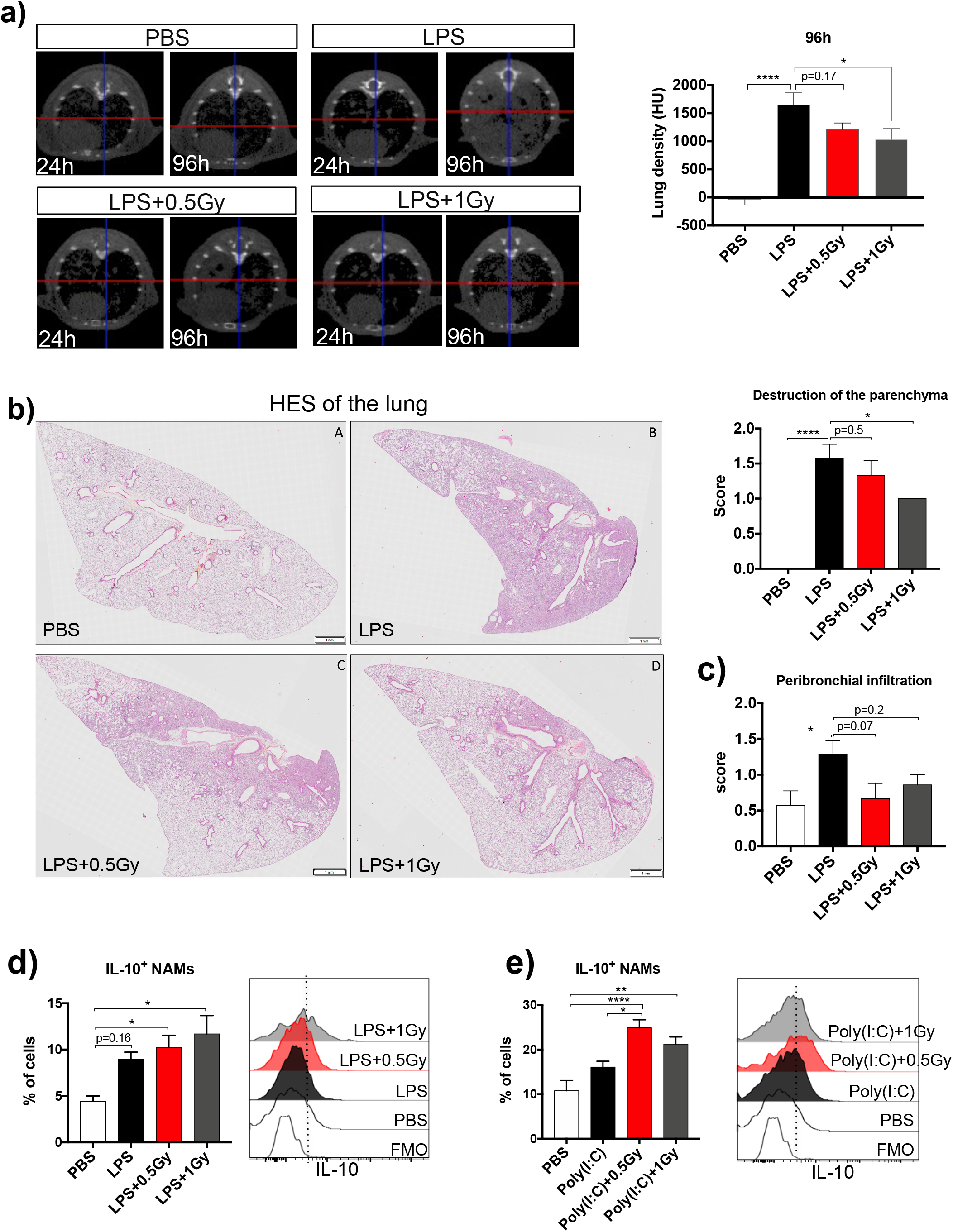
Low doses of RT protect mice from LPS-induced pneumonia and increase IL-10 production by NAMs. Mice were treated with LPS or PBS by intratracheal administration for two consecutive days. Six hours after the second intratracheal administration, mice were irradiated at the whole thorax at 0.5 Gy or 1 Gy. (**A**) Computed tomography (CT) scans of lung density in different treatment groups 24 hours and 96 hours after the first dose of LPS (left) and lung density quantification in the different treatment groups 96 hours after the first dose of LPS. Data are from three independent experiments, n=10-11. (**B**) Representative images (Hematoxylin-eosin-saffran (HES) staining) of lung from LPS treated mice (left, Scale bar=1mm) and scoring of the destruction of the parenchyma (right). (**C**) Scoring of the peribronchial infiltrate. Data are representative of two independent experiments, n=6-7. (**D** and **E**) Eighteen hours after irradiation, the percentages of IL-10^+^ NAMs are presented for each treatment group (left) and representative histograms of the fluorescence associated to IL-10 in NAMs are shown for each treatment group (right). Data were obtained from two independent experiments, n=7-8. For all data, bars indicate mean and error bars indicate ±SEM. *p<0.05, **p<0.01, ***p<0.001, ****p<0.0001 (one-way ANOVA).

NAMs act as a main player to counteract lung inflammation via IL-10 secretion (14). We therefore hypothesized that, in the preclinical LPS pneumonia model, low doses of RT could stimulate IL-10 secretion by NAMs. Our results showed that LPS-induced lung inflammation is associated with a trend towards an increased percentage of NAMs producing IL-10 (Fig. 1D). Interestingly, low doses of RT further increased the percentage of NAMs producing IL-10 compared to the non-irradiated group. Using a Poly(I:C)-induced inflammation model, we confirmed that low doses of RT induced a greater increase of IL-10^+^ NAMs percentage compared to PBS control (Fig. 1E).

### Low doses of RT decrease both histological lung damage and immune cell infiltration during influenza virus infection

To evaluate the effect of low-dose RT on a viral pneumonia model, we treated mice with a lethal dose of PR8 virus (a murine-adapted influenza strain) by intranasal instillation. Two days after the instillation, mice were irradiated at the whole thorax at 0.5, 1Gy or treated with a single dose of dexamethasone. Histopathological analysis of the lung showed that, four days after viral infection, PR8 virus induced a significant increase in both emphysema/alveolar septum rupture and the thickening of the alveolar septum compared to the PBS group (Fig. 2, A and B). Low doses of RT as well as dexamethasone decreased the emphysema/alveolar septum rupture (p=0.0001 for 0.5Gy and p=0.06 for both 1Gy and dexamethasone) and a slight, even if not-significant, reduction of the thickening of the alveolar septum was observed in low RT and dexamethasone groups compared to the PR8 condition. At eight days post-infection, we observed a significant increase in both vascular congestion and red blood cell extravasation in the PR8 group compared to the PBS group (Fig. 2, C and D). Interestingly, low doses of RT or dexamethasone did not exacerbate vascular congestion and red blood cell extravasation compared to the PR8 group. More interestingly, low doses of RT and dexamethasone induced a downward trend in both vascular congestion and red blood cell extravasation compared to the PR8 group.

The analysis of immune cell infiltrate in infected lungs showed that 0.5Gy of RT induced a decreased number of CD45^+^ cells (Fig. 2E) and 1Gy of RT or dexamethasone induced a trend towards a decreased number of CD45^+^ cells. The number of neutrophils, IMs and NAMs significantly increased after PR8 infection, while they were not affected by the low doses of RT or dexamethasone. The number of IMs decreased after low doses of RT (p=0.07 for 1Gy) or dexamethasone treatment (p<0.05). The number of AMs did not change regardless of the treatment. Altogether, our data showed that low doses of RT did not aggravate both tissue damages and inflammatory cell infiltration in the lungs during viral infection.

**Fig. 2.**
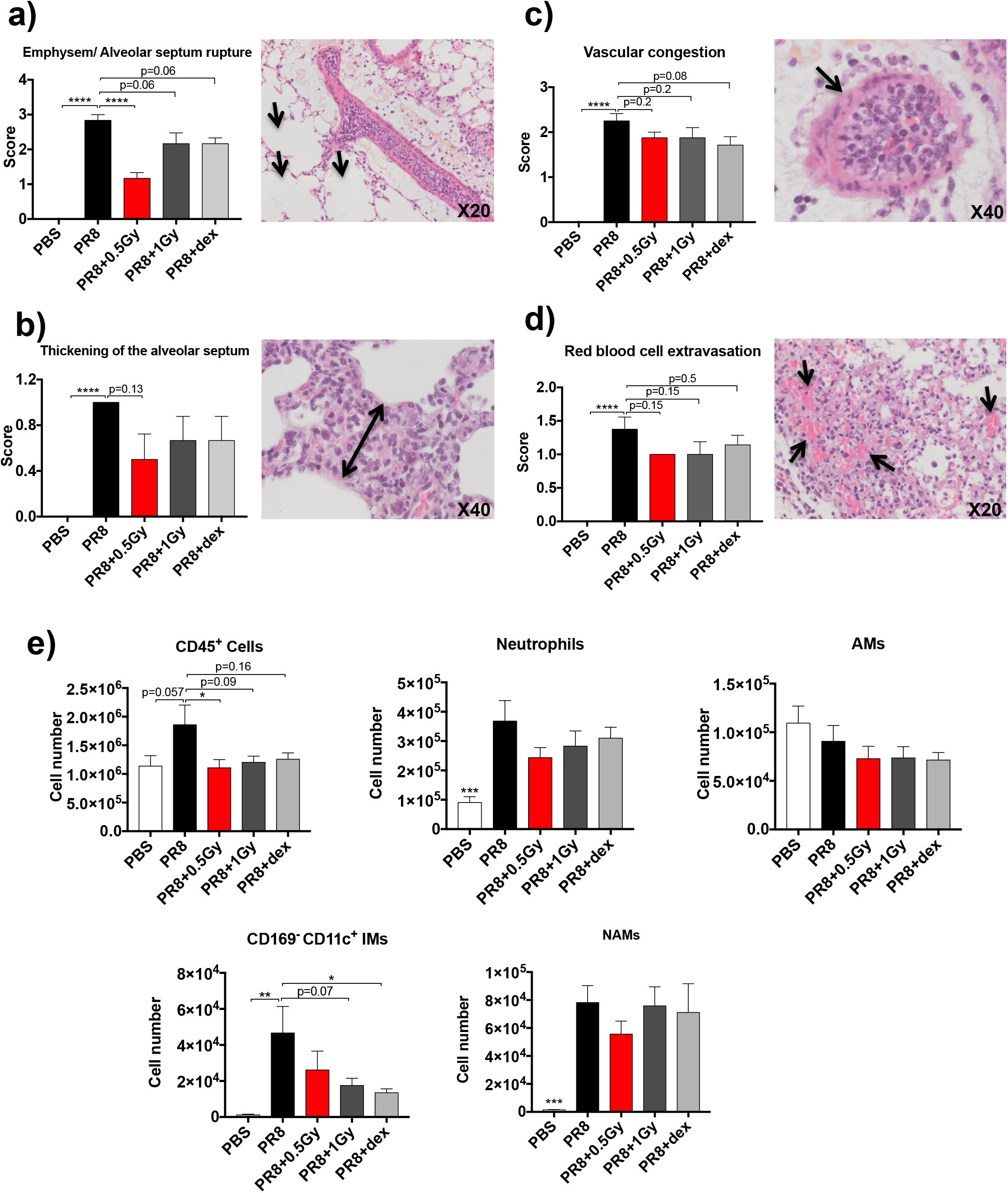
Effect of low-doses RT on both lung tissue damages and immune cell infiltration in H1N1 pneumonia model. Mice were treated with PR8 influenza virus (H1N1) or PBS by intranasale instillation. Two days after, mice were irradiated at the whole thorax at 0.5 Gy or 1 Gy or treated with dexamethasone (dex). (**A**) Emphysem and alveolar septum rupture scoring four days after PR8 infection (left) and representative images (HES staining) of tissue lung from PR8 infected group (right), arrows indicate emphysem. (**B**) Thickening of the alveolar septum scoring four days after PR8 infection (left) and representative images (HES staining) of tissue lung from PR8 infected group (right), double pointed arrow indicates alveolar septum thickening. (**C**) Vascular congestion scoring in the tissue lung eight days after PR8 infection (left) and representative images (HES staining) of tissue lung from PR8 infected group (right), arrow indicate vascular congestion. (**D**) Red blood cell extravasation scoring in the lung tissue eight days after PR8 infection (left) and representative images (HES staining) of lung tissue from PR8 infected group (right), arrows indicate red blood cell extravasation. Data are from two independent experiment, n=6-8. (**E**) Number of CD45+ cells, neutrophils, AMs, CD169^−^ CD11c^+^ IMs and NAMs in murine lungs three days after PR8 infection analyzed by flow cytometry. Data are from two independent experiment, n=6. For all data, bars indicate mean and error bars indicate ±SEM. *p<0.05, **p<0.01, ***p<0.001, ****p<0.0001 (one-way ANOVA).

### Low doses of RT increase IL-10 production by lung immune cells during influenza virus infection

Flow cytometry analysis three days after PR8 infection showed that 0.5Gy of RT, in contrast to dexamethasone treatment, induced a significant increase in the percent of NAMs producing IL-10 (Fig. 3A). The percent of NAMs producing both IFNγand IL-6 significantly decreased after PR8 infection compared to the PBS group, and it was neither affected by low doses of RT nor dexamethasone. Similarly, low doses of RT induced an increase in the percent of both AMs and neutrophils producing IL-10 compared to the other group (Fig. 3, B and C). The percents of AMs and neutrophils producing IFNγand IL-6 were not affected by the different treatments. In agreement with data obtained with LPS and poly(I:C) models, our data clearly show that low doses of RT during viral infection stimulate murine NAMs (as well as alveolar macrophages and neutrophils) to produce IL-10.

**Fig. 3.**
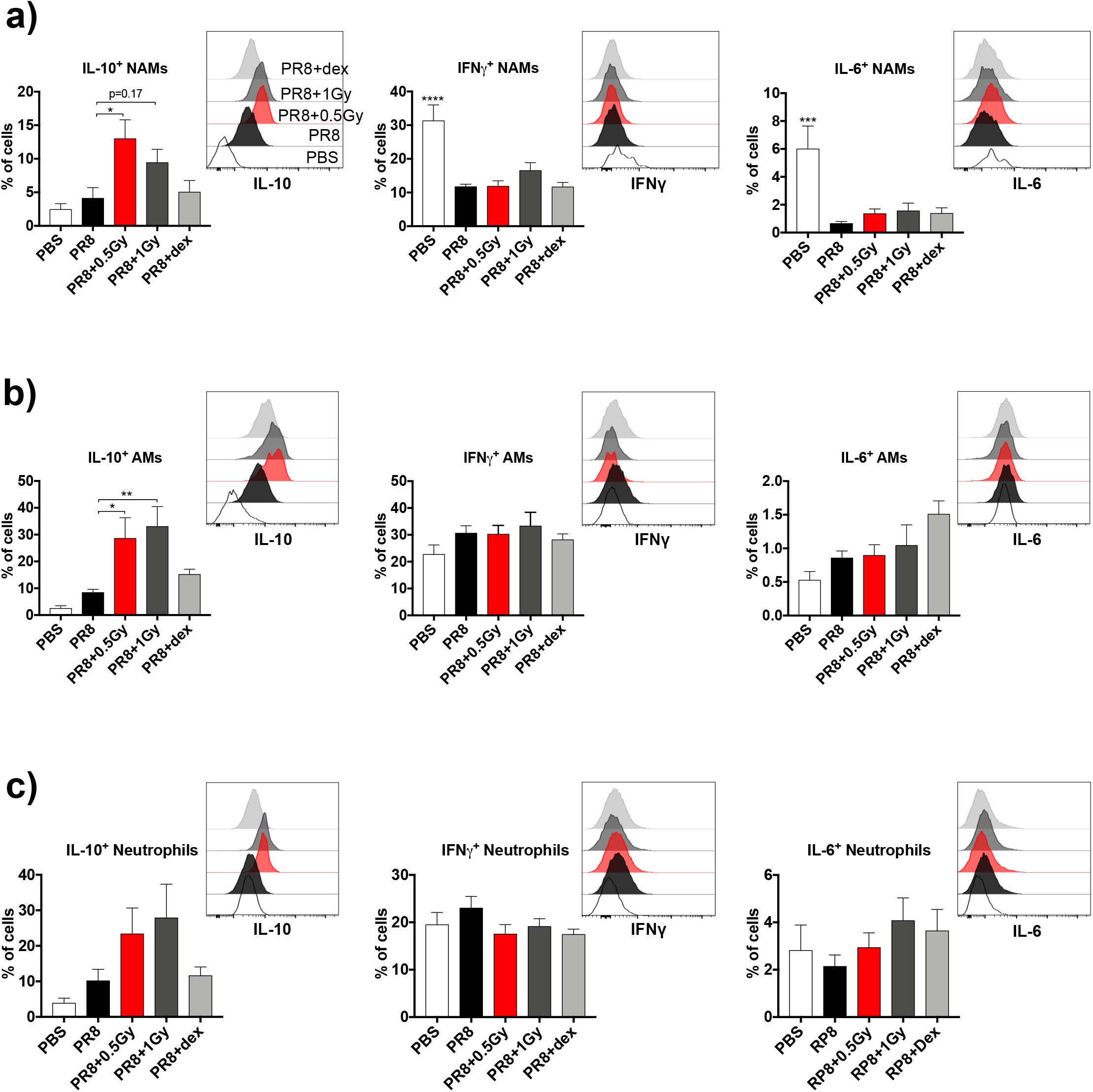
Low doses of RT increase IL-10 production by immune cells in H1N1 pneumonia model. Mice were treated with PR8 influenza virus (H1N1) or PBS by intranasal instillation. Two days later, mice were irradiated at the whole thorax at 0.5 Gy or 1 Gy or treated with dexamethasone (dex). (**A**) Three days after PR8 infection, the percentages of IL-10^+^, IFNγ^+^ and IL-6^+^ NAMs are presented for each treatment group (left) and representative histograms of the mean fluorescence of IL-10, IFNγ, IL-6 in NAMs are shown (right). (**B**) The percentages of IL-10^+^, IFNγ^+^ and IL-6^+^ AMs are presented for each treatment group (left) and representative histograms of the mean fluorescence of IL-10, IFNγ, IL-6 in AMs are shown (right). (**C**) The percentages of IL-10^+^, IFNγ^+^ and IL-6^+^ neutrophils are presented for each treatment group (left) and representative histograms of the mean fluorescence of IL-10, IFNγ, IL-6 in neutrophils are shown (right). Data are from two independent experiments, n=6. For all data, bars indicate mean and error bars indicate ±SEM. *p<0.05, **p<0.01, ***p<0.001, ****p<0.0001 (one-way ANOVA).

### Low doses of RT increase IL-10 production and decrease IFN γ secretion by human lung macrophages *in vitro*

We then aimed to confirm whether low doses of RT could stimulate human lung macrophages to produce IL-10. Sixteen hours after human macrophage irradiation, culture supernatants were analyzed for cytokine secretion and human macrophage activation was analyzed by flow cytometry. The quantification of the supernatants showed that low doses of RT decreased IFNγsecretion and increased IL-10 secretion by Poly(I:C)-stimulated human lung macrophages compared to non-irradiated Poly(I:C)-stimulated ones (Fig. 4A). The levels of IFNα, TNFα, IL-2 and IL-9 were not affected by the low doses of RT.

**Fig. 4.**
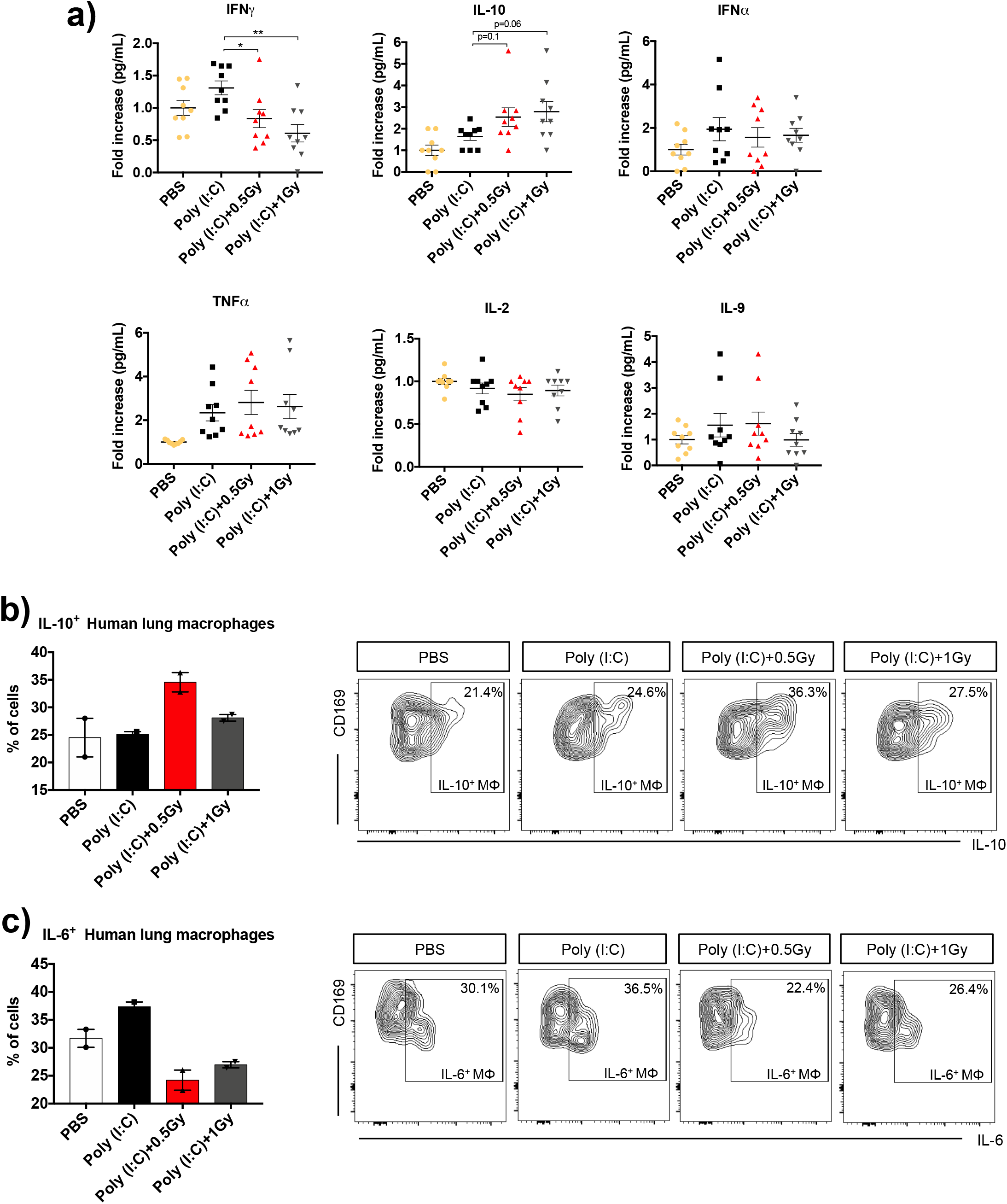
Low doses of RT increase IL-10 production and decrease IFN γ secretion by human lung macrophages *in vitro*. Human lung macrophages were stimulated *in vitro* using the TLR3 ligand polyinosinic:polycytidylic acid Poly(I:C), or treated with PBS (as control). After 6 hours, stimulated macrophages were irradiated at 0.5 Gy or 1 Gy. (A) Sixteen hours after human macrophage irradiation supernatants from cultured macrophages were analyzed for cytokine secretion. Data were obtained from 4 independent experiments, n=9. (**B**) The percentages of IL-10* human macrophages are presented for each treatment group (left) and gating strategy to identify human lung macrophages CD169^+^ producing IL-10 (right). (**C**) The percentages of IL-6^+^ human macrophages are presented for each treatment group (left) and gating strategy to identify human lung macrophages CD169^+^ producing IL-6 (right). For (**B** and **C**) data were obtained from one experiment, n=2. For all data, symbols and bars indicate mean and error bars indicate ±SEM. *p<0.05, **p<0.01 (one-way ANOVA). Data information: human lung macrophages were obtained from healthy lung biopsies.

Subsequently, we performed flow cytometry analysis for human lung macrophages and observed an increase in the percentage of human lung macrophages producing IL-10 after low-dose irradiation of 0.5 Gy compared to other culture conditions (Fig. 4B). Interestingly, the percentage of human lung macrophages producing IL-6 decreased after low-dose irradiation of 0.5 and 1 Gy compared to non-irradiated human lung macrophages (Fig. 4C).

### Low doses of RT had no effect on H1N1 virus expansion in the lungs

Finally, we evaluated the impact of low doses of RT on viral load in the lung. Four days after PR8 infection, broncho-alveolar lavage fluid from infected animals, were analyzed for the measurement of the viral titers. Our results indicated that low doses of RT as well as dexamethasone treatment have no effect on virus levels in the lungs (Fig. 5).

**Fig. 5.**
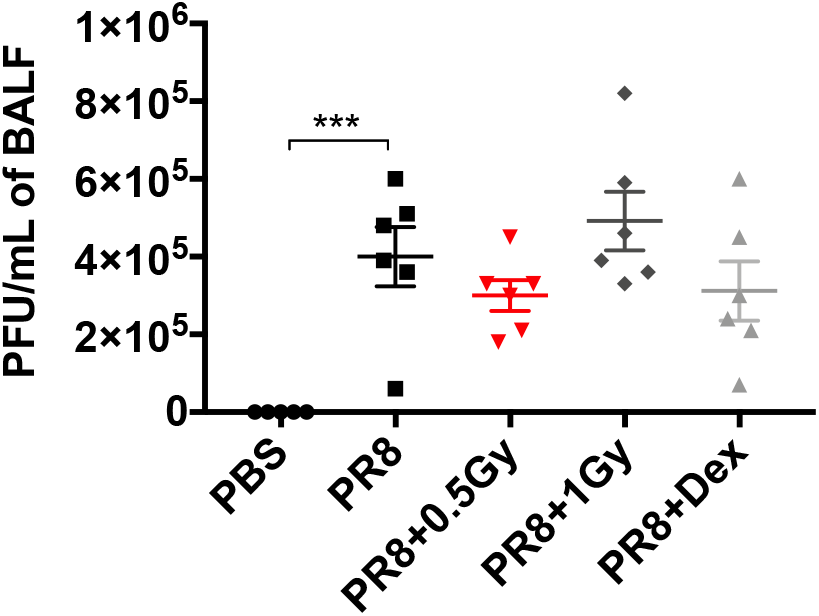
Low doses of RT have no effect on PR8 virus expansion. Mice were treated with PR8 influenza virus (H1N1) or PBS by intra-nasal instillation. Two days later, mice were irradiated at the whole thorax at 0.5 Gy or 1 Gy or treated with dexamethasone (dex). Virus titers were measured in the bronchoalveolar lavage (BAL) four days after animal infection. Data were obtained from two independent experiments, n=6. Symbols indicate mean and error bars indicate ±SEM. ***p<0.001(one-way ANOVA).

## Discussion

In the present study, we described the role of low-dose RT in: 1- protecting mouse lungs from inflammation; 2- further stimulating the production of anti-inflammatory cytokine IL-10 by NAMs *in vivo;* and 3- counterbalancing the effect of the proinflammatory stimuli *in vitro* in human lung macrophages.

The effect of low doses of RT on bone marrow-derived macrophage phenotype and on macrophage cell lines (RAW264.7 and THP-1) has already been reported elsewhere. Interestingly, low dose RT (0.5 Gy) affects the bone marrow-derived macrophage phenotype, however, this strongly depends on the microenvironment (21). Furthermore, a decrease in IL-1 secretion by macrophage cell lines after low doses of RT (0.5 and 0.7Gy) has been reported and LPS-induced TNF production was suppressed in 0.5Gy-irradiated RAW264.7 (22, 23). Accordingly, our results showed that human lung macrophages were affected by low doses of RT (0.5 and 1Gy) *in vitro*. Irradiated human lung macrophages decreased IFNγproduction and increased IL-10 secretion. Furthermore, the percentage of human lung macrophages producing IL-10 increased after low doses of RT, while the percentage of human lung macrophages producing IL-6 decreased after exposure to inflammatory stimuli. Our data demonstrates that low doses of RT are involved in limiting the inflammatory response in favor of an anti-inflammatory response. In preclinical pneumonia models, tissue resident NAMs have been reported to robustly respond to inflammatory stimuli early during infection and to be the main negative regulators of inflammation in the lung *via* IL-10 production as a potential mechanism (14). Similarly, our data confirmed the production of IL-10 by NAMs after Poly(I:C), LPS and PR8 influenza virus infection. Furthermore, we observed that a low dose of RT protects mice from lung inflammation, does not exacerbate lung tissue damages and decreases immune cell infiltration. Whether or not the same NAMs that have been described in mice exist in human lungs with the same immunosuppressive function is not yet defined. Our data from *in vitro*–cultured human lung macrophages are not sufficient to answer this question and further immunohistological/cytofluorimetric analyses of human lung tissue are required.

In pneumonia, several old clinical studies have reported the effectiveness of low doses of RT (5–7). However, after the onset of effective antimicrobial agents, the use of IR in the treatment of patients has been discontinued and the involved mechanism remained unknown. Following the excess death toll related to the COVID-19 pandemic, some radiation oncologists suggested the use of low doses of RT to treat COVID-19^+^ patients suffering from ARDS (24–28), even if this raised some criticism, (29–32) given the uncertainties regarding a potential viral flare-up or an increase in lung tissue damage. In the absence of preclinical data, the scientific community is puzzled between the risks associated with a whole lung irradiation (acute worsening of the patients, and the immediate intrinsic risk of ARDS). Our present study is the first to propose a key role of low doses of RT in the management of pneumonia using preclinical models and suggests lung macrophage reprogramming into immunosuppressive profile as one of the mechanisms involved. Thrombotic events in the lungs of COVID-19 patients following ARDS have been reported (33). Our results showed that PR8 influenza infection induced vessel congestion and obstruction, similar to what is observed in human COVID-19 lung infection. Interestingly, we show that low doses of RT do not worsen this phenomenon of obstructed vessels; on the contrary, there is a downward trend. Furthermore, AMs play a pivotal role in the antiviral defense during influenza virus pneumonia and their depletion induced viral expansion (14). Our results obtained using PR8 influenza pneumonia model indicate that low doses of RT have no effect, neither on the AM population nor on viral expansion, suggesting that the use of low doses of RT would not negatively affect these parameters in viral pneumonias.

Our data show that low doses of RT decrease the tissue damages in the infected mice, and will contribute to define the optimal radiation dose to be used in the clinic, which is the minimal dose to induce a maximal impact on macrophage reprogramming without tissue damages, our data suggest that the 0.5-1Gy range should be considered clinically. In line with our data, several clinical studies including COVID-19 presenting pneumonia patients with advanced age or comorbidities reported an absence of acute toxicity after whole thorax low doses of RT (8, 9). In these clinical studies, the authors reported an improvement in the respiratory function of the patients in the hours following low doses of RT with a marked improvement on the CT scan around 7 days after RT. These observations are in line with the kinetics of our observations. In summary, in line with the recent clinical data, the present study suggests that a single low-dose chest irradiation could be an efficient strategy to resolve human lung inflammation in pneumonia. Low doses of RT induce human lung macrophage reprogramming, in particular the production of the immunosuppressive IL-10 cytokine and the suppression of the inflammatory signals (IFNγ), which could be one of the mechanisms by which low doses of RT protect from human pneumonia (Fig. 6). Our data highlight the effect and good tolerance of low doses of RT on both human and murine macrophage reprogramming and the positive regulation of lung inflammation. Our findings are in agreement with the initial clinical data released from two out of the current 14 activated clinical trials worldwide evaluating chest RT for COVID patients and contribute to justifying the use of RT in this very unusual non-cancer setting. Of note, the radiation doses used empirically in these clinical trials were not uniform; we do believe that carefully defining the lowest as possible effective dose is a major challenge at the heart of the current controversy and our data underscore the fact that doses superior to 1Gy are non-necessary. These data provide the preclinical rational for both use and safety of low doses of RT in situations such as COVID-19-induced ARDS.

**Fig. 6.**
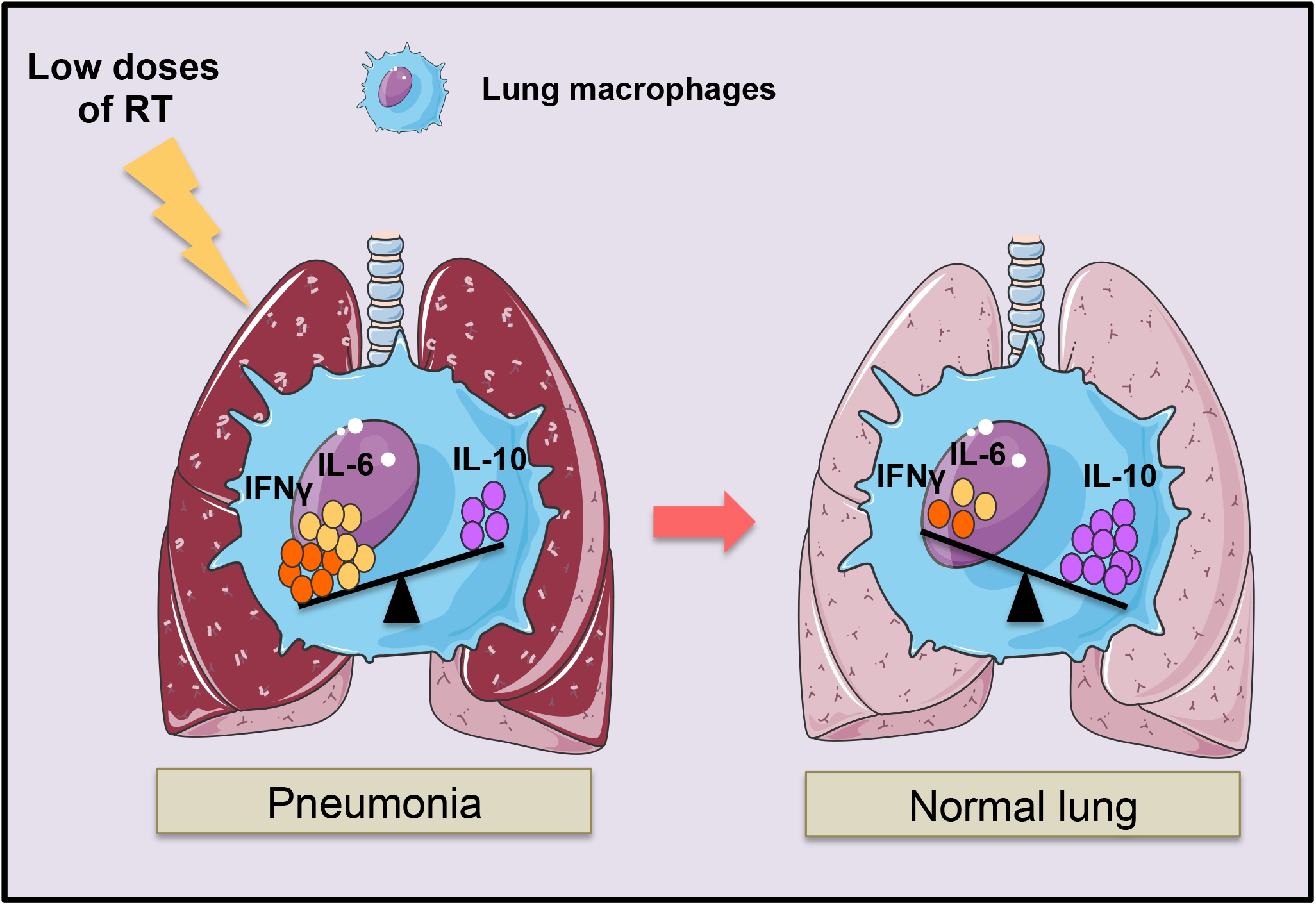
Lung macrophage reprogramming by low doses of RT during pneumonia. Low doses of RT induced the immunosuppressive profile of lung macrophages by increasing the IL-10 production and decreasing the proinflammatory cytokines such as IFNγleading to the lung inflammation resolution.

## Methods

### Human tissue samples

All patients signed an informed consent allowing the use of their surgical specimen for research purposes. The database was declared to the National Board for Informatics and Freedom (Commission Nationale Informatique et Liberté, CNIL, authorization #2217874) and to the National Institute for Health Data (Institut National des Données de Santé, INDS, authorization #MR4316030520). Immediately after anatomical lung resection for lung cancer (n =3) or benign disease (n=1), a 2cm wide peripheral wedge of macroscopically normal lung parenchyma was harvested on the surgical specimen and kept in sterile saline solution at 4°C.

### Animals

Animal procedures were performed according to protocols approved by the Ethical Committee CEEA 26 and in accordance with recommendations for the proper use and care of laboratory animals. For the pneumonia model, female C57BL/6 mice (10 weeks old) were purchased from Janvier Laboratories (France) and housed in the Gustave Roussy animal facility.

### LPS and Poly(I:C) administration

For the pneumonia model, mice were anesthetized (isoflurane), and either LPS (O55:B5) or Poly(I:C) in 50 μl sterile phosphate-buffered saline (PBS), or PBS alone (as control), were administered intratracheally. Mice received two sublethal doses (100 μg and 50 μg) of LPS or (100 μg and 50 μg) Poly(I:C) with a 24-hour rest period between each administration.

### Influenza A virus instillation

Mice received a lethal dose (500PFU) of influenza virus (A/Puerto Rico/8/1934 (H1N1)), in 50 L of PBS by intranasal instillation.

### Irradiation procedure

Six hours after the second administration of LPS or Poly(I:C), the mice were immobilized by anesthesia (2% isoflurane) and locally irradiated at the thorax using a Varian Tube NDI 226 (X-ray machine; 250 kV, tube current: 15 mA, beam filter: 0.2 mm Cu), with a dose rate of 1.08 Gy·min-1. A single dose of 0.5 or 1 Gy was locally delivered to the whole thorax.

### Dexamethasone administration

A single dose of 10mg/kg of dexamethasone (Dexamethasone Mylan 4 mg/1ml, solution for injection, lot n°200147, Mylan) was administered to the mice by intraperitoneal injection.

### Computed tomography (CT) imaging

Mice underwent CT scan at lung level. During scanning, mice were immobilized by anesthesia (2% isoflurane). ImageJ software was used to quantify lung density on axial slices.

### Bronchoalveolar lavage (BAL)

After animal euthanasia, the trachea was cannulated and secured using a silk suture. Cold PBS (500 L) was delivered and retrieved through the cannula. The lavage was repeated three times. BAL fluid was used for virus titration.

### Virus titration

BAL fluids viral titers were determined by standard plaque assay using Madin Darby canine kidney (MDCK) cells. Briefly, confluent monolayers of MDCK cells were infected with serially diluted BAL fluids. Infected monolayers were then covered with semisolid overlay medium (Avicel). 48 hours post-infection, cells were fixed using paraformaldehyde and lysis plaques were counted after crystal violet counterstaining.

### Histopathological analysis and immunohistochemistry

Mouse lungs were fixed in 4% buffered ParaFormaldehye (PFA), paraffin embedded and then cut into 4 m sections. Lung sections were stained with hematoxylin-eosin-saffron (HES) and digitized using a slide scanner (Olympus VS 120).

Histopathological analysis was performed by a Pathologist using a semi-quantitative scoring for the following parameters:

Destruction of the parenchyma: evaluation of the architectural destruction of the lung; Peribronchial infiltrate: presence of inflammatory cells in the peribronchial area; Emphysema and alveolar septum rupture: rupture of the partitions separating the alveoli with the formation of “mega alveoli”; Thickening of the alveolar septum: infiltration of the interalveolar partitions (thickening and / presence of inflammatory cells); Vascular congestion: presence of a large number of vessels filled with immune cells regardless of their caliber; Red blood extravasation: presence of red blood cells outside the vascular lumens, either in the alveoli or in the supporting connective tissue.

### Lung tissue dissociation

Human and mouse lung tissues were digested using the Tumor Dissociation Kit (Miltenyi Biotec) for 30 minutes at 37°C and 1,500 rpm. The cells from the digested lung tissues were filtered using cell strainers (70 μm, Miltenyi Biotech) and used for subsequent experiments.

### Cell culture and irradiation procedure

After washing with PBS and centrifugation (300 g, 4°C, 5 minutes), the human lung cells were suspended and cultured in DMEM-F12 supplemented with both fetal bovine serum (FBS; 10%) and penicillin/streptomycin (1%). Human lung cells were incubated in the indicated medium at 37°C and 5%CO_2_ for 30 minutes. Then, the adherent cells (macrophages and monocytes) were washed using PBS, and the nonadherent cells were discarded. The adherent cells were incubated in fresh medium (DMEM-F12 containing 10% FBS and 1% penicillin/streptomycin) and stimulated with either Poly(I:C) at 1ug/mL or PBS (as control). Six hours after stimulation with the Poly(I:C), macrophages were irradiated using X-RAD320 (X-ray machine; 320 Kev, 4 mA) at a single dose of 0.5 or 1 Gy.

### Flow cytometry

For cultured human lung macrophages staining: anti-CD169 (7-239) and anti-CD11c (REA618, Miltenyi Biotec) were used for membrane staining. Anti-IL-10 (REA842) and anti-IL-6 (REA1037, Miltenyi Biotec) were used for intracellular staining.

For mouse lung cell staining: cell suspensions were incubated with purified anti-mouse CD16/32 (clone 93, BioLegend) for 10 minutes at 4°C. For membrane staining, anti-Ly6G (REA526), anti-CD169 (REA197), anti-CD11c (REA754, Miltenyi Biotec), anti-CD11b (M1/70, BD Horizon™), anti-Ly6C (HK 1.4) and anti-D64 (X54-5/7.1, BioLegend), anti-SiglecF (E50-2440, BD Horizon™) and anti-CD45 () antibodies were used to identify immune cells (CD45^+^), neutrophils (CD45^+^, Ly6G^+^), AMs (CD45^+^, CD11b^−^, CD64^+^, SiglecF^+^, CD11c^+^, CD169^+^), IMs (CD11b^+^ Ly6G^−^ Ly6C^−/low^ CD64^+^, CD11c^+^, CD169^−^) and NAMs (CD11b^+^ Ly6G^−^ Ly6C^−/low^ CD64^+^, CD11c^−^, CD169^+^). Anti-IFNγ(XMG1.2, BD Horizon™), anti-IL-6 (REA1034) and anti-IL-10 (REA1008) were used for intracellular staining. For membrane staining, cells were incubated with the antibody panel at the adapted concentrations for 20 minutes at 4°C. Then, cells were fixed using 4% PFA for 15 minutes at 4°C and permeabilized for intracellular cytokine staining using Perm/Wash Buffer (BD Perm/Wash^TM^). For intracellular staining, cells were pre-activated before membrane staining using Cell Activation Cocktail (with Brefeldin A, Biolegend) for 2 hours at 37°C. Samples were acquired on an LSR Fortessa X20 (BD, Franklin Lakes, NJ) with FACSDiva software, and data were analyzed with FlowJo 10.0.7 software (Tree Star, Inc., Ashland, OR).

### Cytokine analysis

Cytokine concentrations in culture supernatants from *in vitro*-activated human macrophage samples were profiled. The proteins in the supernatant were diluted to 4 mg/mL and analyzed using MACSplex Human cytokine (Miltenyi Biotec) and data were analyzed with FlowLogic 7.3 software.

### Statistical analysis

Statistical analysis was performed using GraphPad Prism 7. One-way ANOVA was used to detect differences among multiple treatment groups. A p value equal to or less than 0.05 was considered significant (*p<0.05, **p<0.01, ***p<0.001, ****p<0.0001). Data are expressed as the mean±standard error of mean (SEM).

## Acknowledgements

The authors thank Patrick Gonin, Karine Ser-Le Roux, Laure Touchard, (PFEP platform), Corinne Laplace-Builhé, Valérie Rouffiac, Philippe Rameau, Yann Lecluse Cyril Catelain (PFIC platform), Olivia Bawa and Hélène Rocheteau (PETRA) at Gustave Roussy for technical assistance.

## Author contributions

LM designed the study, performed the experiments, analyzed the results and wrote the manuscript; CR and SM analyzed CT imaging; PM provided human lung biopsies; MC analyzed histopathological data; BDC and RLG provided the PR8 influenza virus and the measurement of virus titers in the BAL; MM and CC reviewed the manuscript; ED designed and supervised the study and reviewed the manuscript.

